# Erythroid differentiation in mouse erythroleukemia cells is driven via actin filament-tropomodulin3-tropomyosin networks

**DOI:** 10.1101/2021.07.02.448760

**Authors:** Arit Ghosh, Megan Coffin, Richard West, Velia Fowler

## Abstract

Erythroid differentiation (ED) is a complex cellular process entailing morphologically distinct maturation stages of erythroblasts during terminal differentiation. Studies of actin filament assembly and organization during terminal ED have revealed essential roles for the pointed-end actin filament capping proteins, tropomodulins (Tmod1 and Tmod3). Additionally, tropomyosin (Tpm) binding to Tmods is a key feature promoting Tmod-mediated actin filament capping. Global deletion of *Tmod3* leads to embryonic lethality in mice with impaired ED. To test a cell autonomous function for Tmod3 and further decipher its biochemical function during ED, we generated a *Tmod3* knockout in a mouse erythroleukemia cell line (Mel ds19). *Tmod3* knockout cells appeared normal prior to ED, but showed defects during progression of ED, characterized by a marked failure to reduce cell and nuclear size, reduced viability and increased apoptosis. In Mel ds19 cells, both Tpms and actin were preferentially associated with the Triton-X 100 insoluble cytoskeleton during ED, indicating Tpm-coated actin filament assembly during ED. While loss of Tmod3 did not lead to a change in total actin levels, it led to a severe reduction in the proportion of Tpms and actin associated with the Triton-X 100 insoluble cytoskeleton during ED. We conclude that Tmod3-regulation of actin cytoskeleton assembly via Tpms is integral to morphological maturation and cell survival during normal erythroid terminal differentiation.

## Introduction

Definitive erythropoiesis in mammals is characterized by the formation of mature red blood cells (RBCs) in the fetal liver and (postnatally) in the bone marrow via terminal differentiation and maturation of erythroid progenitors, accompanied by assembly of a membrane skeleton framework at the RBC membrane (1). Comprised of ~20 major proteins, the membrane skeleton is a two-dimensional periodic network of α1,β1-spectrin, short actin filaments (F-actin), band 4.1R, ankyrin and other actin-associated proteins (2). The membrane skeleton is critical for maintenance of RBC membrane deformability and stability, and thus cellular integrity, during microcirculation through the vasculature (3). Disruption of membrane skeleton components leads to hemolytic anemias in mice and humans, where RBCs are characterized by abnormal shapes, altered membrane deformability and stability (4, 5). Many hemolytic anemias arise from unstable interactions of spectrin or other network proteins with one another or with the membrane (6), while others may result from impaired membrane skeleton assembly during erythroid differentiation (ED), but this has been less well studied, and the mechanisms of membrane skeleton assembly are not well understood.

The temporal changes in membrane protein expression during ED have been well documented in a variety of systems including mice, chicken, and human RBCs (7, 8). Membrane skeleton proteins such as α1,β1-spectrin, 4.1R and ankyrin are synthesized early in differentiation but are only transiently and inefficiently assembled into the membrane skeleton, with a more stable assembly resulting upon subsequent expression of the anion transporter protein – band3 (7). On the other hand, actin expression decreases during ED (9), with the actin that persists reorganizing from the cytoplasmic cytoskeleton into the membrane skeleton in mature RBCs (10). Correct F-actin assembly is critical for RBC function, since perturbations in G:F (monomeric: filamentous) actin ratio in mature RBCs lead to abnormal membrane deformability in microfluidic assays (11). Notably, RhoGTPases Rac1 and Rac2 and associated signaling molecules that control actin organization are vital for ED, and for correct assembly and maintenance of the RBC membrane skeleton (12). Deletion of some RBC membrane skeleton actin-binding proteins, such as dematin, also affect F-actin assembly into the membrane skeleton and contribute to RBC instability (13). However, the mechanistic details of F-actin regulation in the context of dynamic cytoskeleton remodeling and membrane skeleton assembly during ED remains to be further investigated.

The actin capping proteins, tropomodulins (Tmods), are important players in membrane skeleton organization, ED, and enucleation. Tmods regulate F-actin lengths and stability by capping the slow growing (pointed) filament ends and preventing F-actin assembly and disassembly (14). Tmods’ capping is promoted by binding to Tpms, so that in cells, Tmods predominantly function to stabilize Tpm-coated F-actin in the cytoskeleton (15). The tropomodulin (Tmod) family consists of four members with the ubiquitous Tmod3 expressed early and decreasing during ED, while Tmod1 expression increases during mouse and human ED, similar to other membrane skeleton components, so that Tmod1 is the sole Tmod in mature RBCs (16, 17). While loss of Tmod1 has no effect on membrane skeleton protein expression or assembly during ED (17), a Tmod3 global knockout mouse is embryonic lethal at E14.5-E18.5 with evident anemia at E13.5 due to defective ED of fetal liver definitive erythroblasts with reduced survival, impaired cell cycle exit and reduced enucleation (16). The absence of Tmod3 does not affect Tmod1 expression, implying a unique Tmod3 function during ED that Tmod1 cannot replace. Moreover, enucleating *Tmod3−/−* fetal liver erythroblasts showed aberrant F-actin organization during enucleation (16). Mechanistically, it is not clear how lack of Tmod3 leads to dysregulated ED, and whether Tmod3 regulation of F-actin capping and stability is responsible.

Here, to directly assess the role of Tmod3 in ED and membrane skeleton assembly we employed a murine erythroleukemia cell line (Mel ds19), which has been widely used to investigate many erythroid processes, including ED and membrane skeleton assembly, and is amenable to efficient and rapid genetic manipulation (18, 19). We generated a Tmod3 knockout Mel cell line using CRISPR-Cas9 technology and compared ED induced by DMSO in the parental Mel ds19 cells with Tmod3-knockout Mel cells. Our results demonstrate that lack of *Tmod3* leads to impaired ED with defective assembly of some but not all components of the spectrin-based membrane skeleton. While F-actin, Tmod1 and Tpms all failed to assemble into the Triton-insoluble membrane skeleton, protein 4.1R and a1,β1-spectrin assembly were unaffected in absence of Tmod3. We conclude that Tmod3 is essential for accurate reorganization of F-actin cytoskeleton as a requisite for ED, and that assembly of the major membrane skeleton components, a1,β1-spectrin and protein 4.1R, is not sufficient to sustain normal ED. Additionally, Tmod3-Tpm interactions are critical for Tpm-F-actin assembly, so that Tmod3 functions in concert with Tpms to reorganize and stabilize F-actin into the membrane skeleton during ED.

## Materials and Methods

### Cell culture and erythroid differentiation

Mouse erythroleukemia (Mel) cell line clone ds19 (gift from Dr. Yvette Yien, University of Delaware) were cultured at 37°C (5% CO_2_) in suspension in DMEM media (#10-013-CV), containing 5% fetal bovine serum and 100 IU penicillin, 50 µg/ml streptomycin as described (20). To induce ED, exponentially growing Mel ds19 cells were seeded at a density of 2×10^5^ cells/ml and grown in media supplemented with 2% DMSO for 3 to 5 days (20).

### Generation of *Tmod3* knockout in Mel cells via CRISPR-Cas9

Using the CHOPCHOP web server (http://chopchop.cbu.uib.no/), three different guide RNAs were designed targeting exon 9b (sgRNA1), exon 9a (sgRNA2) and exon 9b (sgRNA3) of the *Tmod3* gene (Primers top/bottom: Table 1). The sgRNA oligo was generated using a primer annealing reaction (sgRNA top 10 µM, sgRNA bottom 10 µM, 1X T4 ligation buffer, 1U T4 PNK) via polynucleotide kinase (PNK) phosphorylation and an annealing program (37°C 30 min, 95°C 5 min; ramp down to 25°C at 5°C per min) as previously described (21). Annealed oligos (diluted 1:200) were cloned into pSp-Cas9(BB)-2A-GFP in a 20 µl reaction – (0.5 µl Cas9-GFP plasmid, 2 µl diluted oligo duplex, 2 µl 10X BbsI buffer, 1 µl 10 mM DTT, 1 µl 10 mM ATP, 1 µl BbsI, 0.5 µl T4 ligase, 12 µl ddH_2_O) using a thermocycler (Biorad T100™) cycles 1-6 (37°C for 5 min, 21°C for 5 min). The resulting product was transformed into DH5α competent cells and propagated for plasmid purification. plasmid was purified using Qiagen Midiprep kit.

**Table 1.**
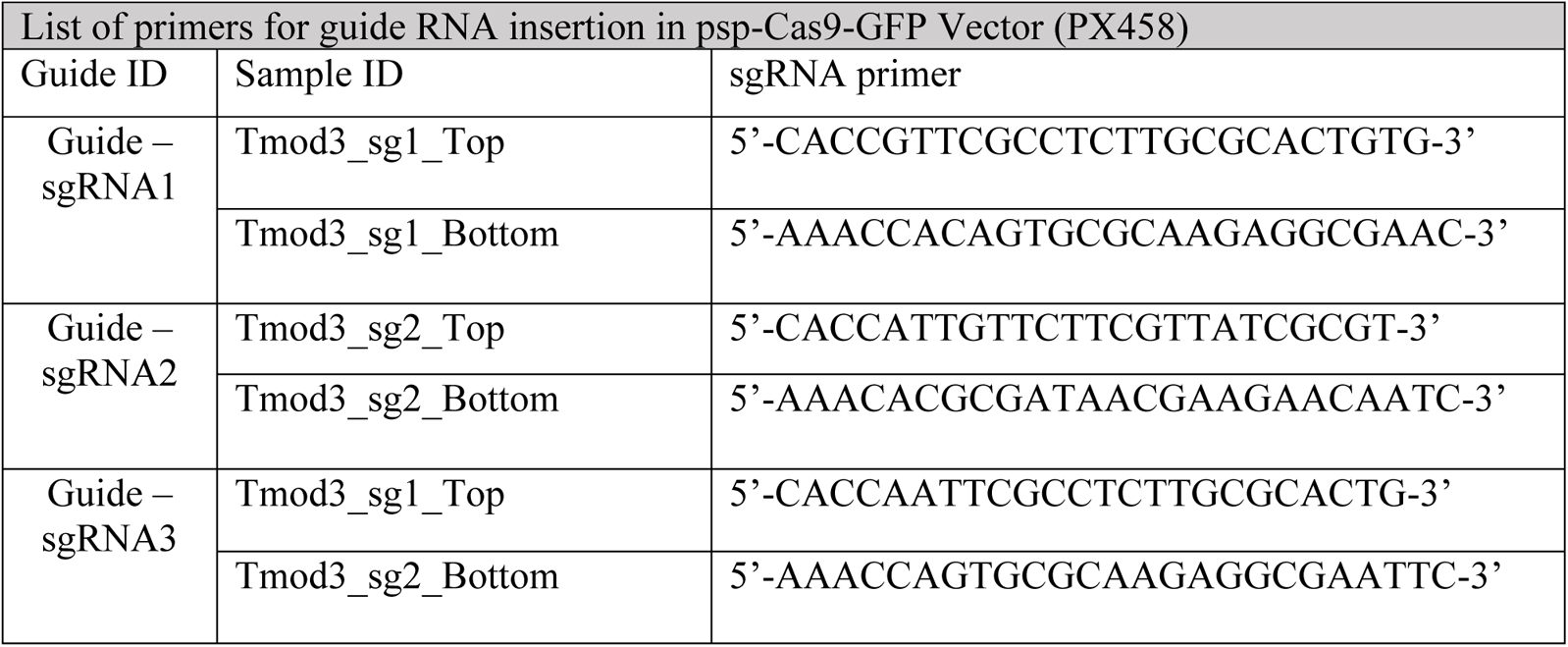

The purified plasmid – pSp-sgRNAx-Cas9(BB)-2A-GFP was electroporated into Mel ds19 cells as previously described (22) with modifications. Cells were washed twice in 1X PBS and resuspended in DMEM (without FBS or antibiotic) and made up to 4×10^6^ cells in 200 µl. 4 µl of 1.5 M NaCl (tissue culture filtered) was added to 200 µl of cells along with 20 µg of the sgTMOD3-Cas9-GFP plasmid (or control plasmid lacking GFP). Cells were transferred to a chilled cuvette (0.4 cm Biorad, #165-2088) and pulse discharged using a setting of 280 V and 975 µF. The cells were transferred to a 10 cm^2^ plate and grown in 10 ml of DMEM (with 5% FBS, 1x Pen/Strep) for 48 hrs after which dead cells were stained with 7AAD (0.05 µg/ml), subjected to flow cytometry in a FACSAria Fusion flow cytometer, and live cells were gated and sorted for GFP expression (Supplemental Figure 1). GFP positive cells were sorted into 96 well plates containing tissue culture filtered spent DMEM media and proliferating colonies were progressively transferred from 48- to 24- to 12-well plates depending on growth. Of the three guide RNAs only sgRNA1 transfection led to viable colonies while the other two guides did not. Several clones (60 from sgRNA1) were screened individually (via western analyses) for the loss of Tmod3 protein. Clone #57 (data not shown) which lacked the Tmod3 protein band of interest was subsequently passaged five times and reanalyzed for the loss of Cas9-GFP from the population with the same gating strategies. Reassessment of Tmod3 protein loss was confirmed via three antibodies specific for various regions of the Tmod3 protein (Figure 1).

**Figure 1.**
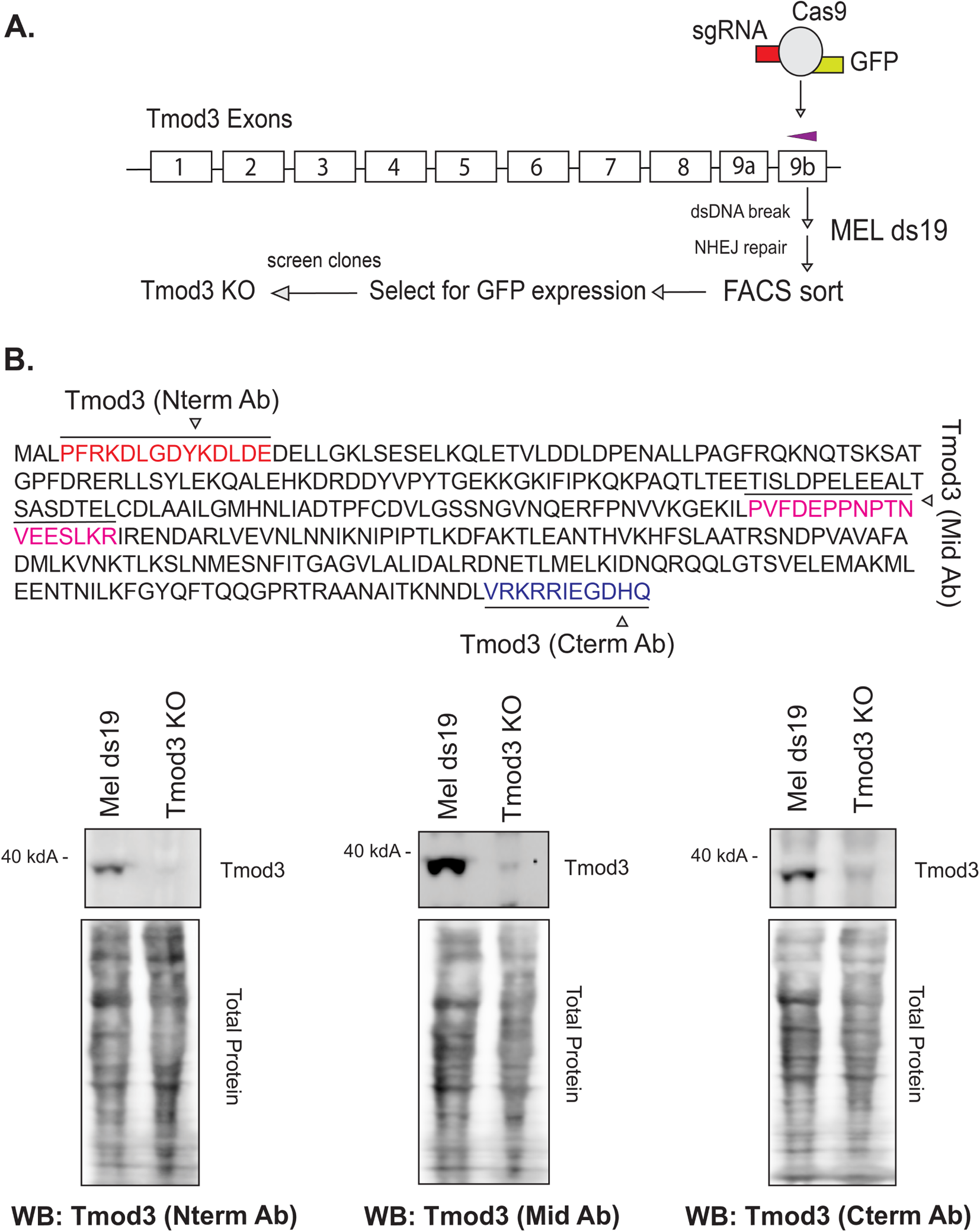
Generation of a *Tmod3* knockout mouse erythroleukemia cell line by CRISPR-Cas9. (A) A *Tmod3* knockout line was created with CRISPR-Cas9 employing a one-plasmid system expressing a guide RNA and Cas9-GFP to target exon 9b on the *Tmod3* gene. Post-electroporation, the cells were sorted for GFP expression via flow cytometry and clones were expanded for screening via western blotting (Supplemental Figure S1). (B) Mouse Tmod3 amino acid sequence with locations of peptides (red – Nterm, blue - Cterm) used to custom prepare chicken anti-Tmod3 antibodies (Genscript). A commercial rabbit anti-Tmod3 antibody (Aviva) raised against the middle region of Tmod3 (pink) was used as well. These three different Tmod3 antibodies detecting various epitopes (Nterm Ab, Mid Ab and Cterm Ab), were used to confirm target protein loss in the *Tmod3* knockout (KO) cells by western blot (upper panels). Total protein was detected using revert stain (LiCoR) (lower panels).

### Cell viability and TMB assay

Cell viability was assessed by Trypan Blue staining using a 1:1 v/v cell to stain ratio and quantified on a BioRad TC20 cell counter to obtain total and live cell counts. Equal numbers of viable cells (2 x 10^5^) were used to carry out the TMB (3,3’,5,5’-tetramethylbenzidine) assay for heme as per manufacturer’s instructions (Pierce #34021). Absorbance was measured at OD 450 nm using a Promega Glomax plate reader.

### Giemsa Staining

Control (untreated), 3d and 5d DMSO treated cells were used for Standard Giemsa staining. 1 x 10^5^ cells were cytospun (1,400 rpm for 3 mins at room temperature; Thermo Cytospin 4) onto glass slides and placed in 100% methanol for 5 mins for fixation. After air drying, the slides were immersed in Giemsa stain (Sigma-Aldrich, GS500), (1:10 diluted in deionized water) and stained for 1 hr. Post-staining, the slides were rinsed in deionized water to remove excess stain and air dried. Slides were examined on a Zeiss AxioImager A2 using a 63X objective lens (numerical aperture, N.A., 1.4) and images acquired with a color camera Axiocam 208, mono camera Axiocam 305 and processed using ZEN 3.0 software. Cell and nuclear area were measured in ImageJ using the freehand selection tool with a paired Student’s T-Test for statistical significance.

### Cellular measurements

*Cell & Nuclear area measurements* – Images from Giemsa staining of control, 3d and 5d cells (Mel ds19 and *Tmod3* KO) were used to calculate cell and nuclear area with the ImageJ freehand selection tool. For each measurement N=30 cells were selected from each condition and respective cell line. Average cell or nuclear area is represented as a horizontal line on dot plots showing all the data points. Experiments were carried out in triplicates. Error bars represent ± SD.

### Flow Cytometry

All cell-based assays for flow cytometric analyses utilized cells initially seeded at 2×10^5^ cells/ml treated with or without 2% DMSO (for differentiation). All experiments were conducted in triplicate and analyzed using FCS Express 7 (Research edition DeNovo Software LLC). Statistical significance testing was done using paired Student’s T-Test. Graphs were generated using Graphpad Prism.

*Annexin V-7AAD staining –* Cells from each condition were centrifuged at 2000 rpm, 2 mins at 4°C, washed with 1X PBS (with 1 mM EDTA) and stained with 5µl Annexin V conjugate in Annexin-binding buffer (23) (Table S2) along with 10µl 7-AAD (dead cell stain) in a final volume of 100µl. The suspension was incubated at room temperature for 15 mins, spun down at 1000 rpm for 3 mins to remove excess stain and resuspended in 500µl of Annexin-binding buffer. Unstained cells were used for gating controls and samples were acquired on a BD FACSAria™ Fusion flow cytometer.

*Cell Event Caspase 3/7 Green detection assay* – CellEvent™ green detection reagent (Thermo #C10423) was used for detection of caspase3/7 activation according to manufacturer’s instructions. Cells were harvested and washed as mentioned above. Detection of activated executioner caspases was done using CellEvent™ reagent according to manufacturer’s instructions. Caspase 3/7 was detected on a BD Accuri C6 flow cytometer in the FITC channel (excitation/emission = 503/530m). Representative flow cytometry panels for percent of populations which are FITC positive [and 7AAD negative (live cells); data not shown] are shown in Figure S3B. The percent of the FITC positive population from each cell line was plotted on a bar graph (Figure 3C) as the mean ± SD for 3d and 5d of growth in DMSO. An unstained sample (Mel ds19) was used as a negative gating control.

**Figure 3.**
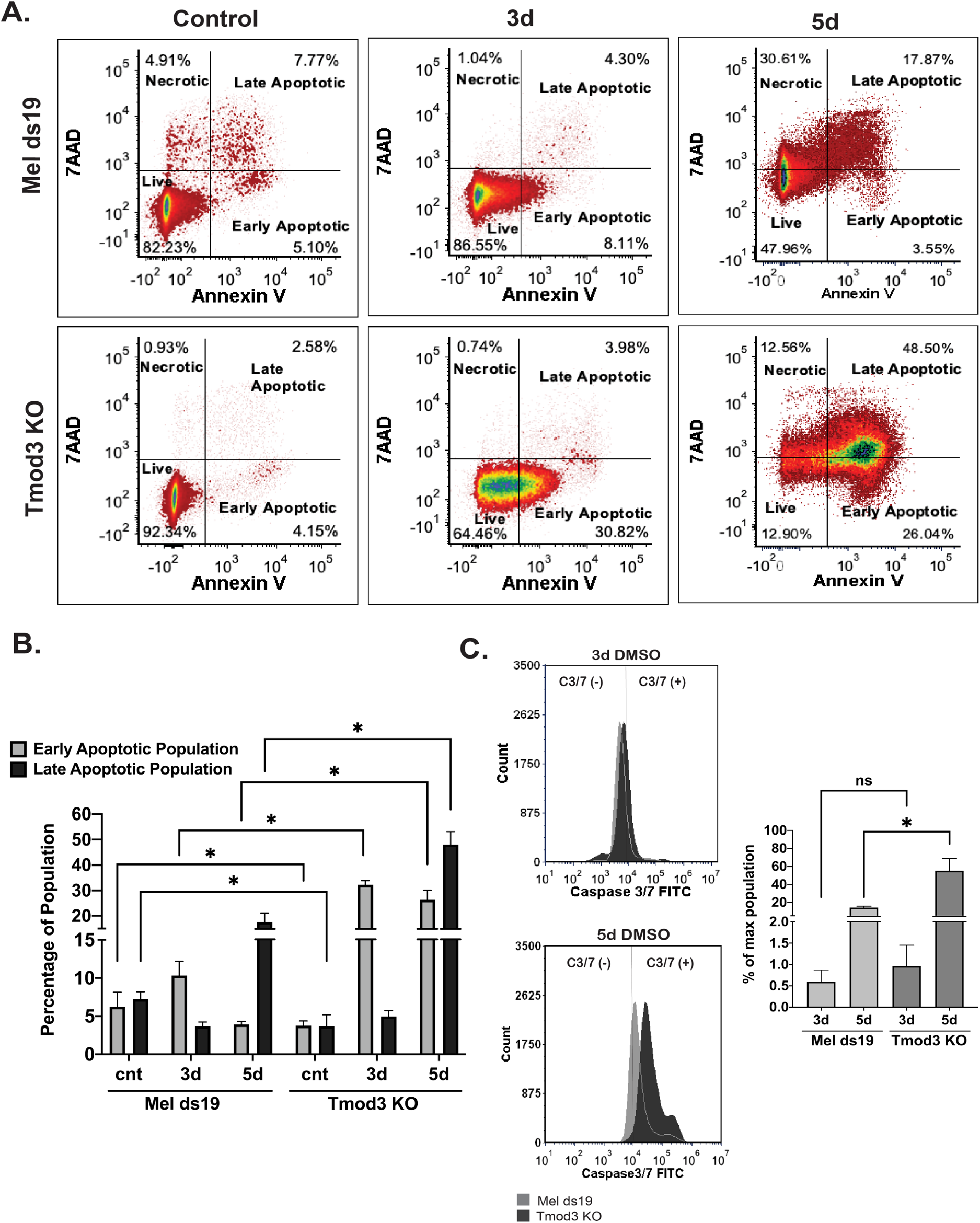
Increased apoptosis during erythroid differentiation in *Tmod3* knockout cells. (A) Annexin V and 7AAD staining in Mel ds19 and *Tmod3* knockout cells was conducted for control (undifferentiated), 3d DMSO and 5d DMSO conditions (differentiating cells). Representative flow cytometry density plots are shown for each timepoint and corresponding cell line. Quadrants denote: live (AnnexinV^low^7AAD^low^), necrotic (AnnexinV^low^7AAD^high^), early apoptotic (AnnexinV^high^7AAD^low^) and late apoptotic (AnnexinV^high^7AAD^high^) cells. (B) Bar graphs showing percentage of cells in early and late apoptotic populations for Mel ds19 and *Tmod3* knockout cells in control (undifferentiated), 3d and 5d DMSO (differentiating) conditions, from flow cytometry data as in A. Values are means ± S.D. (n=3 experiments). *p < 0.05. (C) Caspase 3/7 flow cytometry assays are shown for Mel ds19 and *Tmod3* knockout cells for 3d and 5d timepoints. Left panels, Representative histograms denote Caspase3/7 FITC fluorescence intensities with Mel ds19 in gray and *Tmod3* KO in black. The vertical line on the plots demarcates the negative (C3/7−) and the positive caspase 3/7 signal (C3/7+), based on gating of unstained samples (Supplemental Figure S3A). Right panels, caspase 3/7 positive cells, plotted as a percentage of total population (Caspase 3/7 FITC positive; derived from Supplemental Figure S3B) for control, 3d and 5d cells (Mel ds19 and *Tmod3* KO). Experiments were carried out in triplicate. ns – not significant. *p ≤ 0.05.

*CD71 trafficking –* For determination of surface and internal protein levels, a previously described method was adapted (24) with an Fc antibody (for blocking), CD71 PE (for surface) and CD71 APC (for intracellular). Using the Fix and Perm method (Thermo #GAS003) we were able to detect surface CD71 levels (Figure S2) in the PE channel and internal CD71 levels in the APC channel (post-fixation and permeabilization). Cells were harvested and washed as above. Analyses were carried out on a BD Accuri C6 flow cytometer with appropriate color compensation and unstained controls.

*Propidium iodide staining and cell cycle analysis* – Cell cycle analysis was carried out via Propidium Iodide (PI) staining (control, 3d and 5d) and analyzed on a FACSAria Fusion flow cytometer. Propidium Iodide (Sigma-Aldrich) (1mg/ml) was prepared in a Modified Vindelov buffer (25) [5ml of stock 1mg/ml PI added to 95ml Tris-NaCl (1mM Tris, 1mM NaCl) buffered saline pH 8.0, 70U Ribonuclease A (Sigma-Aldrich), and 0.1 ml IGEPAL CA-630 (Sigma-Aldrich)]. Cells were resuspended in 500µl of PI staining solution and incubated at 4°C for 10 min before analysis. Multicycle analysis on FCS Express 7 software (26) was used to determine diploid G_0_/G_1_, diploid S, diploid G_2_/M phases count vs PE-A (PI stain) histograms (Figure S4).

### Total lysates and western blotting

Total lysates for western blotting were obtained from control, 3d and 5d DMSO samples. Cell pellets (2×10^6^ cells) were incubated in 50 µl of extraction buffer [50 mM Tris-Cl pH 7.4, 150 mM NaCl, 1 mM EDTA, 1.5 mM MgCl_2_, 1% Triton X-100, 1X protease inhibitor cocktail, 1X phosphatase inhibitor (ThermoFisher, Waltham, MA)] for 15 min on ice before sonicating for 10s on ice and centrifuging at 15,000 rpm for 15 mins at 4°C. Protein amount per sample was determined via BCA analysis using BSA as standard (ThermoFisher, Waltham, MA). For total lysates, equal amounts of protein (30 µg) were loaded per lane. For western analyses all samples were electrophoresed on 4-12% SDS (Tris-Glycine) gels (ThermoFisher, Waltham, MA), except for the HbA blot which used 4-20% gels to detect the lower MW (~18 kDA) band for hemoglobin A. Proteins were immunoblotted to PVDF membranes (Millipore Sigma) using the semi-dry transfer (Biorad Trans-Blot Turbo™ #1704150), according to manufacturer’s instructions (LiCoR #926-11010). Total protein was stained by incubating the PVDF membrane in Revert (Li-CoR Biosciences) protein stain and detected in the 700 nm channel with a Biorad Chemidoc Imaging system, prior to antibody labeling. Blocking was conducted using the LiCoR recommended blocking buffer (LiCoR BioSciences#927-70001) and primary antibody incubation was carried out overnight at 4°C with constant shaking in a fresh volume of LiCoR blocking buffer. The following day the blots were washed 3 times with 1X TBST (20mM Tris, 137 mM NaCl, 0.1% Tween-20) and incubated with appropriate secondary antibodies (680RD or 800CW) for 1 hour at RT according to manufacturer’s instructions (https://www.licor.com/bio/applications/quantitative-western-blots/resources) and washed 3 times with 1X TBST before image acquisition. The secondary antibody signals were detected in the 680 nm or 800 nm channels using a Chemidoc. Antibodies are listed in Supplemental Table 1.

### Custom Tmod3 antibody preparation

For preparation of immunopurified Tmod3 antibodies, we utilized Genscript’s (Piscataway, NJ) antibody manufacturing services. The N terminal peptide - PFRKDLGDYKDLDE and a C terminal peptide VRKRRIEGDHQ – in the Tmod3 protein sequence was used to generate chicken anti-Tmod3 polyclonal antibodies. Individuals can contact Dr. Velia M. Fowler, University of Delaware, Newark for these antibodies.

### Triton X-100 subcellular fractionation

For Triton X-100 soluble (S) and insoluble (P) fractions, samples were prepared similar to previous work with modifications (27, 28). Cells from control, 3d and 5d DMSO treatment were spun down at 600g for 10 min at 4°C and washed twice with 1X PBS to remove cell debris and excess media. Samples were incubated in Buffer A, an actin stabilization buffer [0.1 M PIPES pH 6.9, 30% Glycerol, 5% DMSO, 2 mM MgCl_2_, 100 mM KCl, 1 mM EGTA, 1% Triton X-100, 1 mM ATP, 1X protease inhibitor cocktail (ThermoFisher, Waltham, MA)] for 10 min on ice and centrifuged at 100,000 g for 45 mins at 4°C. The supernatant was removed as the Triton X-100 soluble (S) fraction and an equivalent amount of Buffer B, actin depolymerization buffer [0.1 M PIPES pH 6.9, 2 mM MgCl_2_, 100 mM CaCl_2_, 1X protease inhibitor cocktail, 5 µM Cytochalasin D (ThermoFisher, Waltham, MA)] was added to resuspend the pellet (P). This Triton X-100 insoluble fraction (P) was incubated with Buffer B for 30 min on ice and then sonicated for 10s to completely disperse and resuspend the pellet. The S and P samples were solubilized by addition of an equal volume of 2X Laemmli SDS-PAGE sample buffer (Biorad) and boiling for 5 min. Equal volumes of each (S and P) sample were loaded on 4-12% SDS-PAGE gels, and western analysis was conducted as above.

### Western blot quantification

*Western blot (total protein) analyses* – Band intensity was calculated using ImageJ and normalized to Total-Protein Revert™ staining before antibody labeling. Total protein lanes as detected from Revert™ staining were quantified to estimate the standard protein for normalization for each western blot. For each quantification, total area under the curve from image quantification was converted into percent using gel analysis options tool in ImageJ as described (http://www.navbo.info/DensitometricAnalysys-NIHimage.pdf). Percent peak values for each protein (western blot) were normalized to total protein values. The average of the arbitrary units as ratiometric values of percent protein (western blot) to percent protein (total protein) are shown as bar graphs in Fig 4A and Fig 6A.

**Figure 4.**
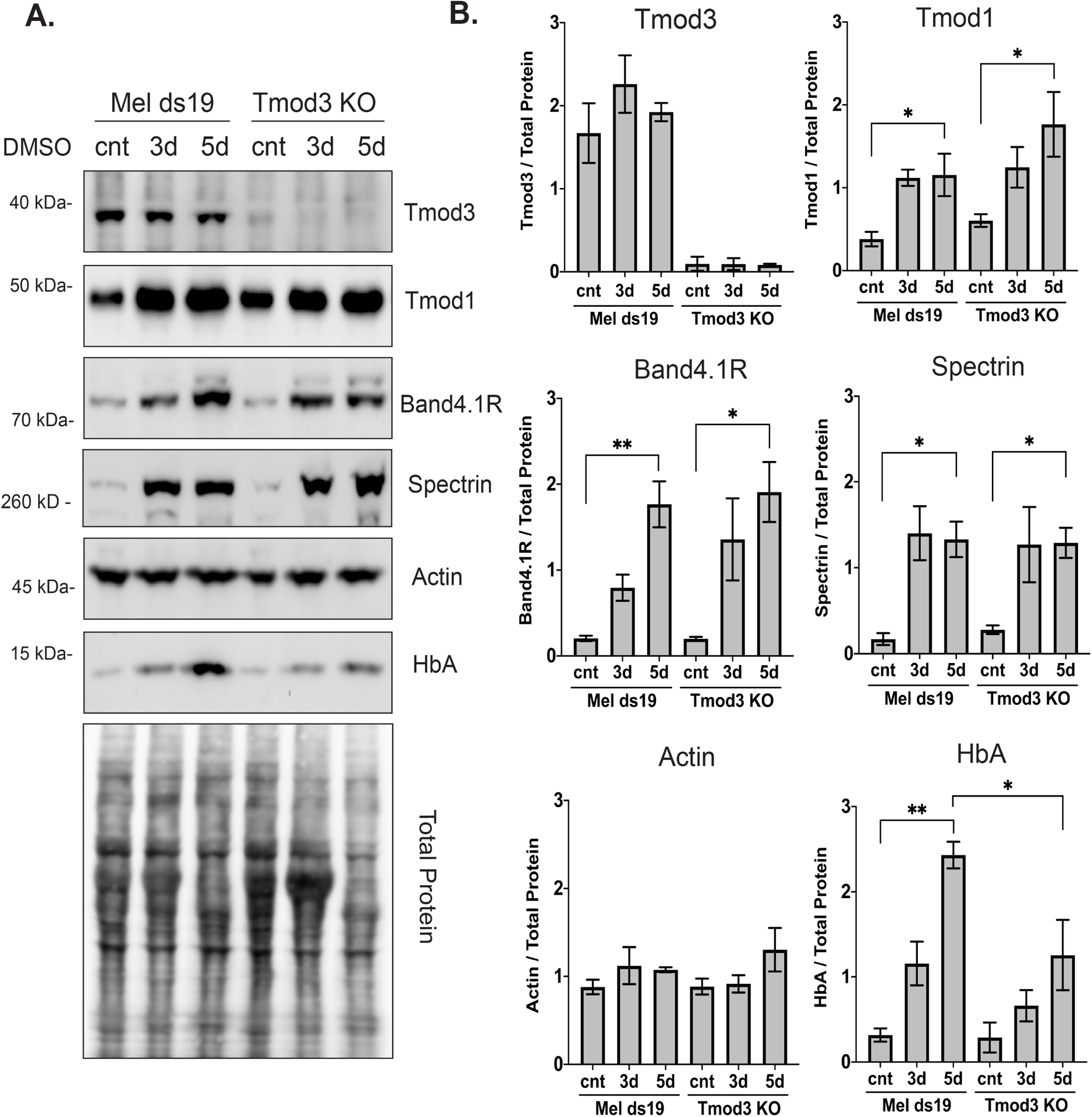
Loss of *Tmod3* does not affect increases in levels of Band 4.1R, a1,β1-Spectrin, Tmod1 and Hemoglobin A (HbA) during erythroid differentiation. (A) Immunoblots of Tmod3, Tmod1, Band 4.1R, a1,β1-Spectrin, Actin and HbA in Mel ds19 and *Tmod3* knockout cells grown in absence of DMSO for 5d (control) or in presence of DMSO for 3d and 5d. Each lane was loaded with 30 μg of total lysates from respective timepoints for each cell line and immunoblotted for proteins as indicated. Bottom panel, Total protein is shown as Revert™ 700 protein stain and used as a loading control. Representative immunoblots are shown. (B) Bar graphs showing relative levels of protein of interest, normalized to total protein, quantified from immunoblots, and denoted as arbitrary units on Y axes for comparison. Values represent means ± SD (N=3 independent experiments). Tmod1, band 4.1R, a1,β1-Spectrin and HbA levels increase during ED in Mel ds19 and *Tmod3* KO cells, while Tmod3 and actin levels remain constant. No Tmod3 is detected in the KO cells. * p ≤ 0.05, ** p ≤ 0.01

**Figure 6.**
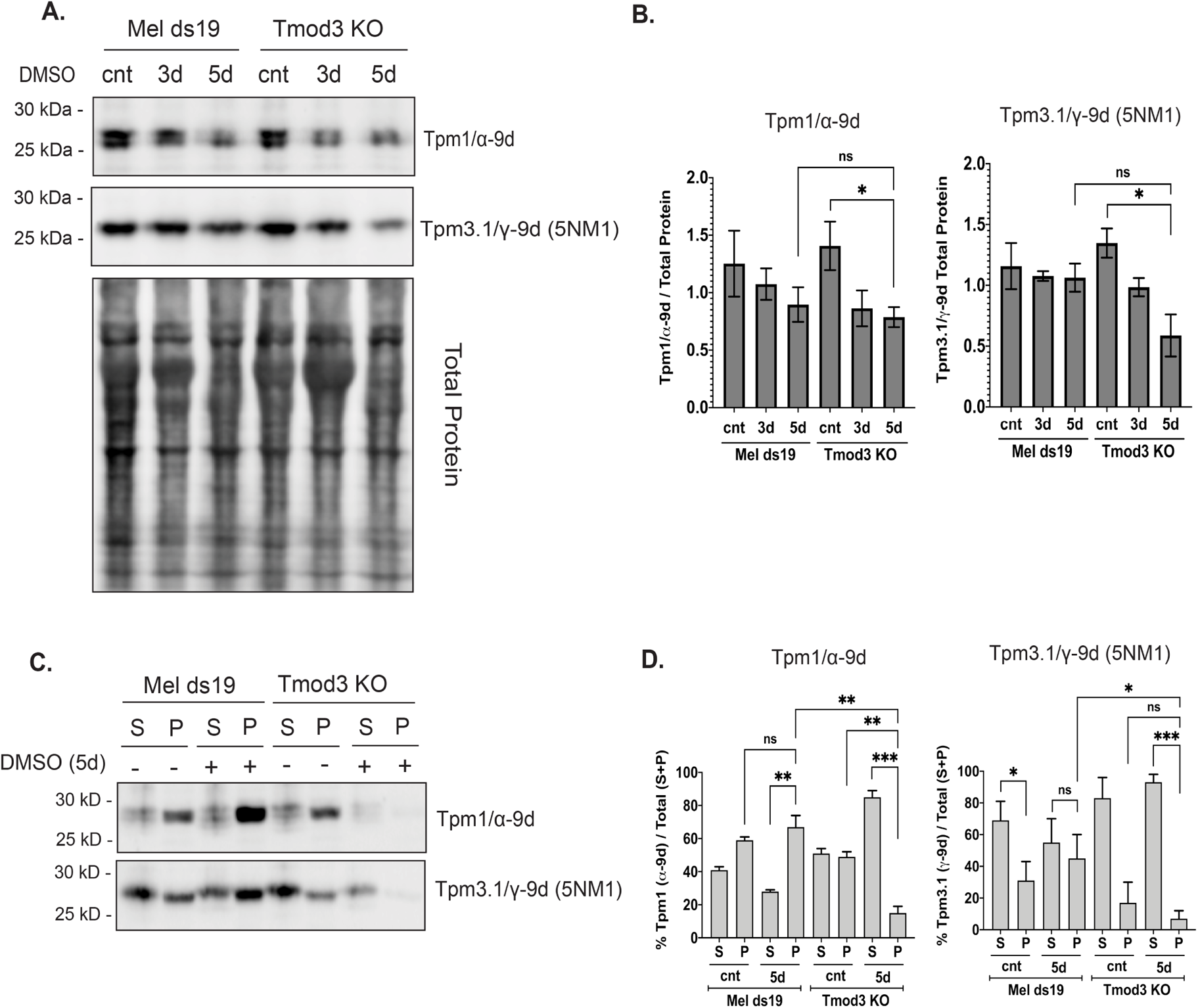
Tropomyosin levels and assembly into the Triton-insoluble membrane skeleton are reduced during erythroid differentiation of *Tmod3* knockout cells. (A) Immunoblots of tropomyosins, Tpm1/α-9d and Tpm3.1/γ-9d (5NM1), in Mel ds19 and *Tmod3* KO cells during ED via immunoblotting. Total protein (revert stain, LiCor) was used as loading control. (B) Bar graphs showing relative levels of Tpms, normalized to total protein, quantified from blots as in A, denoted as arbitrary units on the Y axes. Tpm1/α-9d and Tpm3.1/γ-9d (5NM1) did not increase during ED in Mel ds19 cells, while loss of Tmod3 leads to significant decrease in both Tpms during differentiation. (C) Immunoblots of Tpm1/α-9d and Tpm3.1/γ-9d (5NM1) in Triton X-100 soluble (S) and insoluble (P) fractions from Mel ds19 and *Tmod3* knockout cells grown in the absence (−) or presence (+) of DMSO for 5d cells. Each lane was loaded with equivalent volumes of S and P (10 μl), and representative immunoblots are shown. Total protein is shown as revert protein staining (LiCoR). (D) Percent of Tpm1/α-9d or Tpm3.1/γ-9d (5NM1) in S and P fractions expressed as a percentage of total S+P. In Mel ds19 cells, Tpm1/α-9d levels consistently associate with P fractions in both control and differentiated cells, while Tpm3.1/γ-9d (5NM1) redistributes from S to P fraction as differentiation proceeds. Lack of Tmod3 led to complete loss of both Tpm1/α-9d and Tpm3.1/γ-9d (5NM1) from the P fractions during differentiation. Values represent means ± SD. N=3 independent experiments. ns – not significant, * p ≤ 0.05, ** p ≤ 0.01, *** p ≤ 0.001.

*Western blot (S and P fraction) analyses* – Band intensities and percent values from the western analyses for respective fractions S or P, were measured as described above using ImageJ, and percent values for S and P fractions were added to obtained total intensity. The intensity of S or P fraction was divided by total S+P and represented as an average ± SD (N=3) for each timepoint (control, 3d and 5d) for the respective cell lines.

### Statistical analysis

Data shown in dot plots and bar graphs are mean ± standard deviation (SD). Differences between means were detected using Student’s Paired T-tests. Statistical analysis was performed using GraphPad Prism 7.03 software. Statistical significance was defined as \**p* ≤ 0.05, \*\**p* ≤ 0.01, \*\*\**p* ≤ 0.001, \*\*\*\**p* < 0.0001.

## Results

### Generation of a *Tmod3* knockout in Mel cells

We showed previously that Tmod3 is required for terminal differentiation of mouse fetal liver definitive erythroblasts *in vivo*, but whether this was a consequence of Tmod3 regulation of F-actin was not explored (16). To directly examine the role of Tmod3 in ED, we adopted a CRISPR-Cas9 mediated knockout methodology to perturb the *Tmod3* gene in a mouse erythroleukemia cell line, Mel ds19 (19). We directed disruption of *Tmod3* gene exon 9b via a sgRNA-Cas9-GFP construct leading to a double-strand break followed by canonical non-homology end joining (NHEJ) directed repair (Figure 1A). The cells were sorted for GFP-Cas9 expression and viable clones were expanded and screened for loss of Tmod3 via western blotting (Figure S1). To confirm the identity of the derived clone as a true Tmod3 knockout we utilized antibodies detecting epitopes located in the N-terminal and C-terminal regions of Tmod3 (custom-made antibodies) as well as a commercial antibody detecting an epitope in the middle region of the Tmod3 protein (Supplemental Table 1, Materials and Methods). Lack of Tmod3 in the selected clone confirmed that our clone of interest was a bonafide Tmod3 knockout cell line (Figure 1B). The removal of Cas9-GFP was also tested in the selected clone via passaging and retesting for GFP expression using flow cytometry (Figure S1).

### Lack of *Tmod3* leads to impaired Mel cell differentiation

To characterize the effects of *Tmod3* gene deletion, we analyzed cell viability (using trypan blue staining) during differentiation induced by growth in media containing 2% DMSO. The Mel ds19 control cells varied between 88-90% live cells after 5d in DMSO, with a cell concentration of ~6×10^6^ cells/ml. In contrast, the Tmod3 KO cells had a ~1.4-fold decrease in viability (~65% live cells) after 5d in DMSO with a total cell count of ~2.5×10^6^ cells/ml (Figure 2A). To assess the efficacy of DMSO induction of *in vitro* ED we investigated total hemoglobin production via a chromogenic assay utilizing TMB (3,3’,5,5’-tetramethylbenzidine) as the substrate. As expected, DMSO treated Mel ds19 cells showed a 2-fold increase in TMB signal by 3d, which was sustained at day 5, signifying accumulation of hemoglobin during ED. While the Tmod3 knockout cells showed an increase in hemoglobin signal at 3d, there was a significant decrease at 5d compared to the control Mel ds19 cells, suggesting lack of sustained expression of hemoglobin during ED in absence of Tmod3 (Figure 2B). When we looked at surface and internal transferrin receptor (CD71) levels, we found that Tmod3 KO cells showed a decrease in surface CD71 similar to Mel ds19 cells (Figure S2), indicating that Tmod3 KO cells had some normal features of ED even as overall differentiation was impaired.

**Figure 2.**
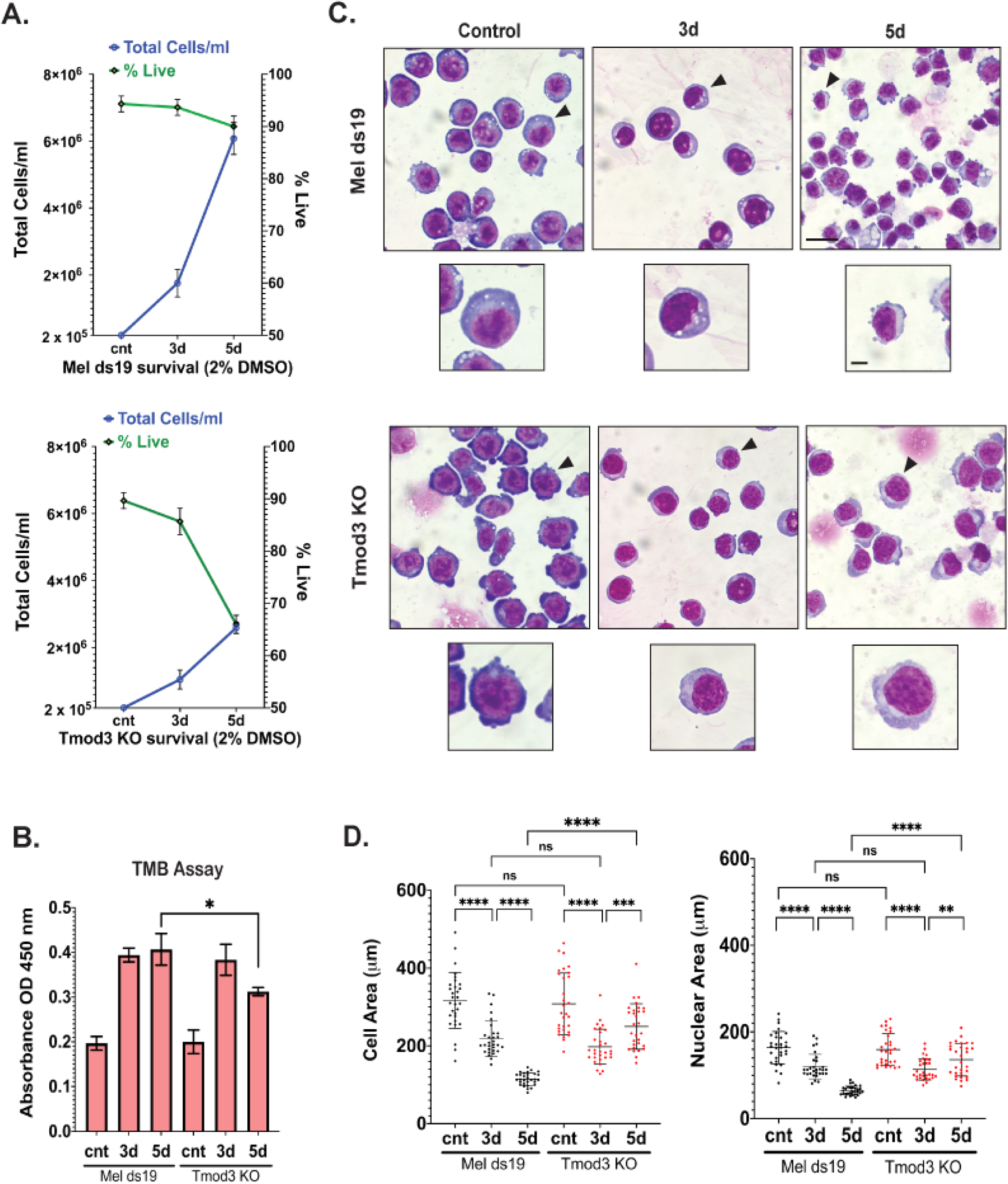
*Tmod3* knockout leads to decreased cell viability, total hemoglobin and impaired erythroid differentiation. (A) Mel ds19 and *Tmod3* KO cells were seeded at 2 x 10^5^ cells/ml and grown for 3-5 days in culture media supplemented with 2% DMSO to induce erythroid differentiation (ED). The control timepoint denotes undifferentiated cells after 5 days of culture without DMSO treatment. At each timepoint cells were stained with trypan blue to ascertain cell viability and the percentage of live cells was counted. (B) The TMB (3,3’,5,5’-tetramethylbenzidine) assay was used to determine total hemoglobin content in Mel ds19 and *Tmod3* KO cell lines, using equal number (2×10^5^) of live cells (trypan blue staining) for each time point (control, 3d and 5d). The graph represents an average of triplicates with a mean ± standard deviation. (C) Wright-Giemsa staining was used to analyze morphological changes in the two cell lines during ED. Representative images are shown for control, 3d and 5d DMSO conditions. Insets display a single cell at a higher magnification from the main panel (indicated with an arrowhead) to illustrate morphological differences between the two cell lines. Scale bar = 10 μm. (D). Scatter plots representing mean cell and nuclear area for Mel ds19 and *Tmod3* KO cells were obtained from Giemsa staining (N=30 cells; data shown is representative of three independent experiments). Horizontal line on scatter plots represents average cell or nuclear area, with SD indicated. Cell Area (µM) – Mel ds19 (control – 316.6 ± 71.6, 3d – 218.3 ± 45.7, 5d – 114.6 ± 15.9), *Tmod3* KO (control – 308.0 ± 79.3, 3d – 197.7 ± 45.0, 5d – 249.9 ± 58.8). Nuclear Area (µM) – Mel ds19 (control – 163.8 ± 37.0, 3d – 120.1 ± 28.2, 5d – 64.7 ± 10.2), *Tmod3* KO (control – 159.4 ± 36.6, 3d – 113.5 ± 24.6, 5d – 136.1 ± 37.2). ns – not significant, ** p ≤ 0.01, *** p ≤ 0.001, **** p ≤ 0.0001.

We performed Wright-Giemsa staining to visualize morphological changes in the differentiating cells at 3d and 5d DMSO (Figure 2C). As control Mel ds19 cells differentiate, they show a gradual decrease in both cell and nuclear area, from 0d to 3d to 5d of DMSO treatment, as expected (Figure 2C-D) (29). Before differentiation, the cells display a deep blue cytoplasm and a prominent nucleus. After 3d the cells are smaller with a reduced cytoplasmic area and lighter colored cytoplasm similar to polychromatic erythroblasts (30) (Figures 2C-D). At 5d, the cell and nuclear area of the control Mel ds19 cells decreased further (Figure 2C-D). While Tmod3 KO cells resembled Mel ds19 cells in appearance with decreased cell and nuclear area after 3d, suggesting initial progression of ED, the KO cells failed to show any further cell and nuclear area decreases at 5d. Instead, the sizes of the 5d KO cells were increased compared to 3d KO cells, with a considerably larger cell (~2.5 fold) and nuclear (~3 fold) area as compared to 5d Mel ds19 cells (Figure 2D). Such a pattern is reminiscent of a similar trend for total hemoglobin production which increased at 3d but reversed and decreased at 5d for the KO, as shown above (TMB assay, Figure 2B). This suggests a temporal role for Tmod3 protein in the later stages of terminal ED, and that Tmod3 function is important for the transition from basophilic to polychromatic erythroblast.

### Apoptosis is elevated in absence of *Tmod3* during ED

Since a major decrease in viability was observed for *Tmod3* KO Mel cells during ED, we next looked for changes in apoptosis in the KO cells compared to the parent ds19 cells. Using Annexin V (AV) as a readout for apoptosis and 7AAD as a stain for dead cells, we looked at live, necrotic, early, and late apoptotic populations during ED with flow cytometry (Figure 3A) (23). During proliferation of undifferentiated cells in absence of DMSO, apoptotic populations were low for both parent Mel ds19 cells and *Tmod3* KO cells, with the latter having a ~3 fold lower late apoptotic population compared to parental cells, whereas early apoptotic populations showed no significant differences between the two cell lines (Figure 3A, 3B). At 3d of differentiation, the *Tmod3* KO showed a significantly higher degree of early apoptotic cells (~30%, ~3.5 fold more) compared to Mel ds19, while late apoptotic populations were similar. At 5d, the *Tmod3 KO* had an even greater proportion of both early (~28%) and late (~48%) apoptotic cell populations compared to Mel ds19 cells, in which only ~4% of cells were early and ~18% late apoptotic cells (Figure 3A, 3B). When we looked at cell cycle parameters for 5d of ED, both ds19 and KO cells appeared arrested at the G1 stage (Figure S4) with the *Tmod3* KO showing only ~5% increased cellular granularity for the G1 (2N) population compared to Mel ds19, indicative of more internal complexity (i.e. less differentiated) in the KO.

Executioner caspases 3/7 are integral for proper ED, with increased caspase3/7 required during terminal differentiation (31). Therefore, to further assess the abnormal increase in apoptosis in the *Tmod3* KO cells, we conducted caspase 3/7 assays via flow cytometry for undifferentiated (control), 3d and 5d timepoints. This showed that *Tmod3* KO cells (at 5d DMSO) had ~4-fold higher levels of activated caspases compared to Mel ds19 cells during ED. Based on the observed increases in early and late apoptotic cell populations (AV^high^7AAD^high^) in the *Tmod3* KO cells, it is possible that overactivation of caspases in *Tmod3* KO during ED precipitates cells towards cell death.

### Loss of *Tmod3* reduces Tmod1 and actin assembly into the membrane skeleton

Tmod3 capping of Tpm-coated F-actin promotes assembly and stability of actin cytoskeletal structures such as the Tpm3.1-F-actin-spectrin network on lateral membranes of polarized epithelial cells (32), and the Tpm3.1-g-actin network in sarcoplasmic reticulum of skeletal muscle (33). A hallmark of terminal ED is the increased expression and assembly of the spectrin-F-actin filament network of the membrane skeleton, containing protein 4.1R, α1,β1-spectrin, Tmod1, and Tpms, among other components, despite a decrease in total actin levels (8). To examine whether loss of Tmod3 affected membrane skeleton protein levels and assembly in Mel cells, we measured total levels of actin, α1,β1-spectrin, protein 4.1R and Tmod1 and their assembly into the Triton-X-100 insoluble membrane skeleton during ED. Western analyses of total lysates (control, 3d and 5d) revealed that as differentiation proceeds Mel ds19 cells show ~1.5 to 2-fold increases in α1,β1-spectrin, protein 4.1R and Tmod1 levels, similar to primary murine erythroblasts (8) while total actin levels remained the same (Figure 4A, 4B), Loss of Tmod3, however, did not affect the increase in total levels of α1,β1-spectrin, protein 4.1R, Tmod1 or actin, which were all unchanged. Hemoglobin levels (HbA) also increased during ED in Mel ds19 cells, but this was reduced in *Tmod3* KO cells, as shown above by the TMB assay (Figure 2). Thus, expression of membrane skeleton components appears normal despite the absence of Tmod3, with reduced HbA expression and impaired morphological ED and cell death shown above.

To examine whether membrane skeleton assembly was affected by loss of Tmod3, we carried out sub-cellular fractionation using Triton X-100 for soluble (S) pools and insoluble pellet (P) fractions containing F-actin and the membrane skeleton (Figure 5A). First, we investigated membrane skeleton assembly of actin, α1/β1-spectrin, protein 4.1R and Tmod1 in the Mel ds19 cell line containing Tmod3. In undifferentiated Mel ds19 cells, nearly ~70% of actin was in the S fraction, with the remaining ~30% in the P fraction (Figure 5). Induction of ED led to redistribution of actin pools so that about 50% of the actin was now in the pellet, indicating that ED is accompanied by F-actin polymerization and assembly into the membrane skeleton, as reported previously (34). Similar to actin, Tmod1 also redistributed from the S to the P fraction during ED in Mel ds19, indicating Tmod1 assembly into the membrane skeleton in control cells. By contrast, Tmod3 was highly abundant in the S fractions in both undifferentiated cells and at 5d ED (~80-90%), with only a small proportion associating with the P fraction during ED (~5% in P fraction at 0d and ~10% at 5d of ED). Consistent with membrane skeleton assembly during ED of Mel ds19 cells, protein 4.1R was enriched in the P fraction (35). On the other hand, α1,β1-spectrin was found in both S and P fractions during ED similar to previous studies showing α1,β-spectrin polypeptides accumulate in excess during differentiation (35, 36).

**Figure 5.**
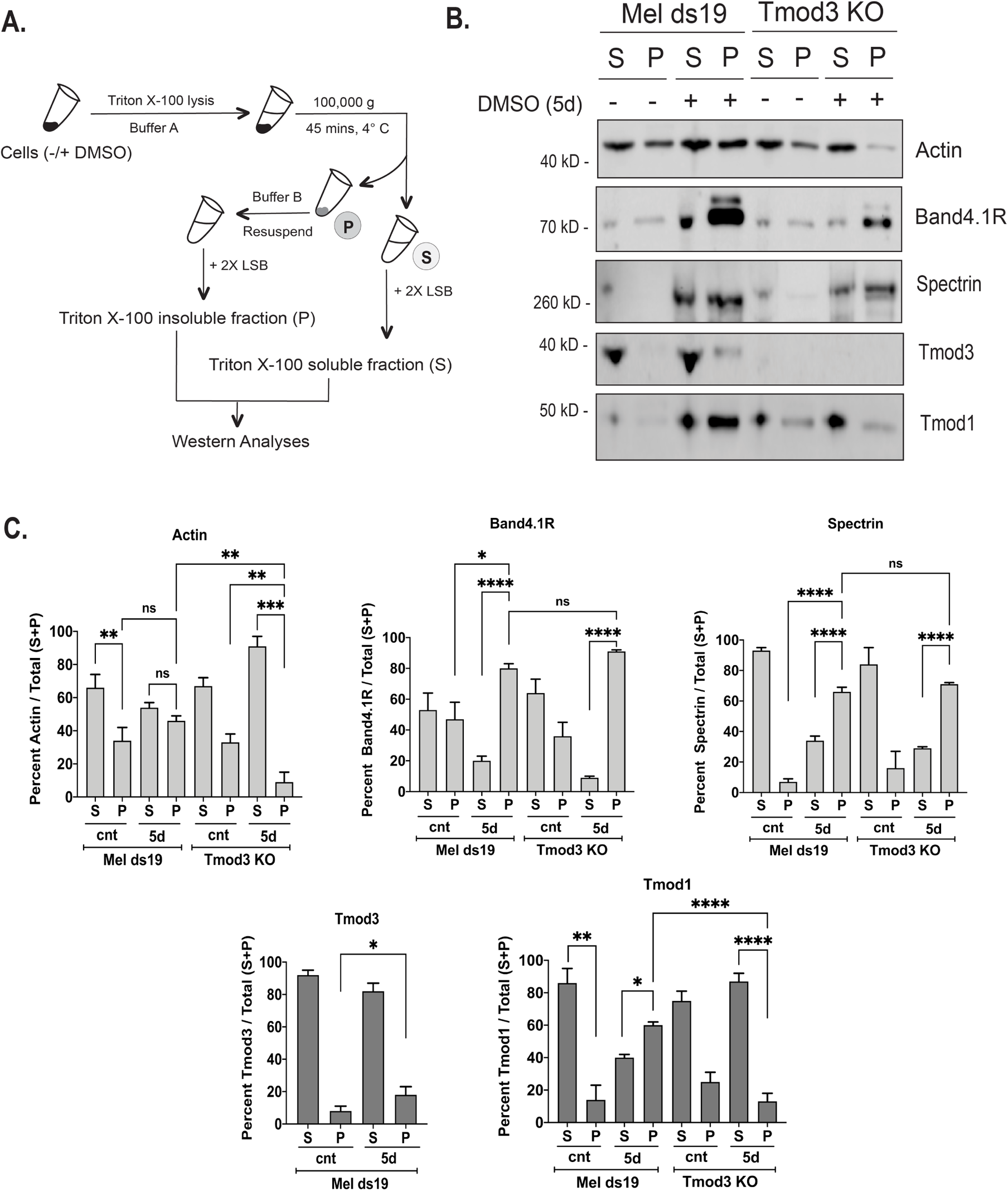
Loss of Tmod3 leads to reduced assembly of actin and Tmod1, but not protein 4.1R or a1,β1-Spectrin, into the Triton-insoluble membrane skeleton during erythroid differentiation. (A) Experimental workflow for actin fractionation into Triton X-100 soluble (S) and insoluble (P) fractions. Cells growing with or without DMSO were lysed in Triton X-100 based Buffer A and the supernatant was removed as the Triton X-100 soluble fraction (S) and the resulting pellet was resolubilized in Buffer B (actin depolymerizing buffer). Equal volumes of S and P fractions were used for western analyses. (B) Immunoblots of actin, band 4.1R, a1,β1-Spectrin, Tmod1 and Tmod3 in Triton X-100 soluble (S) and insoluble (P) fractions from Mel ds19 or *Tmod3* KO cells grown in the absence (−) or presence (+) of DMSO for 5d. Equivalent volumes of S and P fractions (10 μl) were loaded in each lane (see Materials and Methods), and representative immunoblots are shown. (B) Percent in S and P fractions expressed as a percentage of the total S+P. Values for actin, Band4.1R, a1,β1-Spectrin, Tmod3 and Tmod1 in S and P fractions compared to total S+P are shown as means ± SD. Values represent N=3 independent experiments. ns – not significant, * p ≤ 0.05, ** p ≤ 0.01, *** p ≤ 0.001, **** p ≤ 0.0001.

Loss of Tmod3 had an adverse effect on actin as well as Tmod1 distribution during ED (Figure 5A, 5B). While undifferentiated KO cells showed a similar distribution of actin and Tmod1 in the S and P fractions as compared to Mel ds19 cells, lack of Tmod3 during ED led to reduced assembly of both actin and Tmod1 into the P fractions at 5d. During ED, the relative proportion of actin and Tmod1 in the P fraction in the Tmod3 KO was ~4-5 fold lower than for the Mel ds19 cells. Even so, band 4.1R and α1/β1-spectrin showed normal assembly into the TX-100 insoluble membrane skeleton in the Tmod3 KO during ED, similar to parental Mel ds19 cells. This indicates that Tmod1 and actin assembly into the membrane skeleton depend upon Tmod3, while α1,β1-spectrin and protein 4.1R can assemble into the membrane skeleton via an actin and Tmod-independent pathway.

### Tropomyosin assembly during ED is dysregulated in absence of Tmod3

Tropomyosins (Tpm) are considered master regulators of the actin cytoskeleton, binding along F-actin to promote filament stability and regulating interactions with other actin binding proteins (37). Mouse and human RBCs contain two Tpm isoforms, Tpm1.9 and Tpm3.1, that are associated with the short F-actin in the membrane skeleton (38). To determine which Tpm proteins were expressed in Mel ds19 and *Tmod3* KO Mel cells, we used a series of antibodies specific for exon α-9d (Tpm1), exon γ-9d (Tpm3.1), and exon δ-9d (Tpm4) to screen Mel ds19 cell extracts (data not shown) (39). Based on these results, we selected two Tpms, Tpm1 (α-9d) and Tpm 3.1 (γ-9d, 5NM1), for further study during ED. Similar to actin and Tmod3 (Figure 4), Tpm1 and Tpm3.1/5NM1 levels do not increase during ED (Figure 6A), unlike Tmod1, protein 4.1R and a1.β1-spectrin, whose levels increase significantly during ED (Figure 4). In Tmod3 KO cells, levels of both Tpm1 and Tpm3.1 decreased significantly during ED, by about 50% and 60%, respectively (Figure 6A, B), again unlike Tmod1, protein 4.1R and a1,β1-spectrin whose expression levels increased even in the absence of Tmod3 (Figure 4).

To assess Tpm assembly with F-actin into the membrane skeleton during ED, we examined the proportion of Tpms in the Triton X-100 soluble (S) and insoluble (P) fractions (40) (Figure 6C, D). In Mel ds19 undifferentiated cells, Tpm1 was preferentially associated with the P fraction, a distribution maintained during differentiation, while Tpm3.1 was ~2 fold higher in the S vs P fraction in undifferentiated cells, redistributing to a ~1:1 ratio during ED (Figure 6C, 6D). By contrast, in Tmod3 KO cells, both Tpms were predominantly enriched in the S fractions. Tpm1 was evenly distributed between S and P fractions in undifferentiated cells, and failed to associate with P fractions upon ED, so that Tpm1 levels were >4 fold higher in the S compared to P fractions. The preferential enrichment of Tpm in the S fraction of Tmod3 KO cells was more marked for Tpm3.1, which was ~4 fold higher in S vs P fractions even in undifferentiated cells and did not change during ED. Failure of these Tpms to associate with the P fractions in the differentiated Tmod3 KO cells correlates to lack of F-actin assembly (P fractions) in the Tmod3 KO as shown above (Figure 5). Thus, loss of Tmod3 results in the inability to assemble Tpm1- or Tpm3.1-coated F-actin during ED.

## Discussion

In this study, we investigated the role of the pointed-end actin capping protein, Tmod3, in erythroid differentiation (ED) and F-actin assembly. Our results show that loss of Tmod3 in mouse erythroleukemia cells leads to impaired ED in Mel cells, based on decreased accumulation of hemoglobin, failure to reduce cell and nuclear size, reduced viability and increased apoptosis. These results parallel impaired ED observed for definitive ED in fetal liver of *Tmod3−/−* mice in our previous study (16). While *Tmod3−/−* fetal liver erythroid cells also exhibit impaired cell cycle exit, reduced enucleation frequency and abnormal F-actin during enucleation, parental Mel ds19 cells do not normally enucleate, and loss of Tmod3 does not affect cell cycle progression during ED in Mel cells. Thus, Tmod3’s role in cell cycle exit in mouse fetal liver may be restricted to late stages of ED and/or be an indirect consequence of absence of Tmod3 in supporting cells such as macrophages (16). On the other hand, our results showing that loss of Tmod3 leads to impaired ED in Mel cells indicate a cell autonomous function for Tmod3 in ED prior to enucleation.

Tmod3 is required for correct F-actin assembly during ED. *Tmod3−/−* Mel ds19 cells showed decreased assembly of Tmod1, Tpms and F-actin into the Triton-insoluble membrane skeleton during ED. Moreover, protein levels of Tpm1/α-9d and Tpm3.1/γ-9d (5NM1), but not Tmod1 or F-actin, are greatly reduced in *Tmod3−/−* Mel ds19 cells during ED. Since Tmod3 promotes Tpms binding to F-actin, stabilizing Tpm-coated F-actin (14, 15, 38), loss of Tmod3 may lead to Tpms’ dissociation and F-actin instability. This could preclude subsequent recruitment of Tmod1 to F-actin for correct membrane skeleton assembly as ED progresses. In contrast to a Tmod3 requirement for assembly of Tmod1, Tpms and F-actin during ED, loss of Tmod3 is without effect on assembly of α1,β1-spectrin or protein 4.1R into the Triton-insoluble membrane skeleton. This suggests that α1,β1-spectrin and 4.1R assemble independently of F-actin, likely binding to the membrane via ankyrin or protein 4.1R linkages to Band3. Thus, Tmod3 regulation of Tpm-Tmod1 mediated F-actin assembly is integral to cell survival and progression of ED, but not to membrane skeleton assembly, and dysregulated F-actin assembly may explain the abnormal F-actin organization and impaired enucleation in *Tmod3−/−* fetal liver erythroblasts (16).

Tmod3 is not present in mature RBCs, where Tmod1 caps the pointed ends of the short Tpm-coated actin filaments in the spectrin-F-actin lattice (17). Consistent with this, Tmod1 expression increases dramatically during ED in Mel ds19 cells, similar to primary mouse and human erythroblasts (16) and shows a ~40% increased association with the Triton-insoluble fraction, whereas Tmod3 levels do not increase, nor does it assemble into the Triton-insoluble fraction during ED. It is likely that increased Tmod1 during ED competes with Tmod3 for binding to pointed ends of F-actin assembling into the membrane skeleton, precluding association of Tmod3. This is supported by appearance of Tmod3 in *Tmod1−/−* mouse RBCs, with Tmod3 now assembling into the membrane skeleton (17). By contrast, loss of Tmod3 in Mel ds19 cells is not compensated by increased expression or assembly of Tmod1, and instead dysregulates Tmod1 association with the actin cytoskeleton during ED. Thus, Tmod3 is essential for Tmod1 assembly but not vice versa. Moreover, absence of Tmod1 does not affect ED, since *Tmod1−/−* mice have a mild compensated spheroelliptocytic anemia with increased reticulocytosis (17) and normal distributions of differentiating erythroblasts in fetal liver and bone marrow (41). Thus, despite not being a component of the final membrane skeleton, Tmod3 has a critical function in assembly of a key subset of membrane skeletal proteins – Tmod1, Tpms and F-actin.

In addition to Tmod3, other actin regulatory proteins that are not components of the erythrocyte membrane skeleton have also been shown to be critical for ED. For example, lack of Rac1 and Rac2 in mice alters actin assembly in RBCs causing membrane skeleton disruption, microcytic anemia and reticulocytosis (42). Rac1/2 KO mouse RBC ghosts have increased levels of actin and adducin phosphorylation, but Tmod1, Tpms and other proteins are unaffected. Actin and adducin associations with the Triton-insoluble membrane skeleton are also reduced, while Tmod1 and Tpm are unaffected, suggesting Rac1/2 regulation of actin assembly during ED via a divergent pathway from that controlled by Tmod3, which controls Tmod1 and Tpm assembly (12, 42). Integrity of the actin cytoskeleton in Mel ds19 cells has also been reported to depend on actin polymerization stimulated by WASp which activates the Arp2/3 complex (43). A Was knockout in Mel cells has no change in total actin but an altered ratio of monomeric G-actin to polymeric F-actin. Lack of Hem-1, a WAVE complex member which activates Arp2/3 in hematopoietic cells, leads to abnormal erythrocyte shapes and aberrant F-actin foci in *Hem-1−/−* mouse RBCs (44). Additionally, *Hem-1−/−* RBC ghosts display reduced levels of membrane skeletal proteins such as α1,β1-spectrin, adducin, dematin, ankyrin, Band3, Band4.1R, and Tmod1, as well as increased adducin phosphorylation. This suggests an important role for actin regulation during ED by proteins that do not become part of the final membrane skeleton, as we report here for Tmod3 in Mel ds19 cells.

Our studies in Mel cells and fetal liver erythroblasts have both shown that loss of Tmod3 leads to reduced survival due to elevated apoptosis with increase in caspase3/7 activity. How does the erythroid cell relay the signals from identification of a destabilized membrane to cellular machineries that result in apoptosis? Several studies have demonstrated a role for the actin cytoskeleton as a trigger for apoptosis (45). Treatment of HeLa cells with cytochalasin B (inhibitor of actin polymerization) leads to caspase-mediated cytochrome c release indicative of actin-dependent mitochondrial membrane destability (46). Gelsolin (barbed end F-actin binding protein) can induce apoptosis by dissociating the G-actin:DNAase I complex, leading to nuclear localization and activation of DNAse I (47). Bcl-2 proteins have also been shown to have a direct link to actin-mediated apoptosis as the pro-survival Bcl-X_L_ protein suppresses apoptosis induced via cytochalasin D in Jurkat cells (48). Moreover, increased Bcl-X_L_ expression is required for survival of differentiating erythroid cells (49). One can speculate that loss of an actin capping protein such as Tmod3 might act as a trigger for activation of signaling cascades leading to abrogation of increase in Bcl-X_L_ expression, culminating in erythroid cell death. Further studies to elucidate Tmod3’s binding partners in erythroblasts will be necessary to elucidate the mechanistic connections between Tmod3 regulation of actin assembly with cell survival and progression of ED.

## Acknowledgments

We thank Dr. Yvette Yien for the gift of the Mel ds19 cell line. We also thank Dr. Kasturi Pal and Mr. Declan Cole for assistance in clonal selection of the Tmod3 knockout cells.

## Author contributions

A.Ghosh – performed and designed experiments, analyzed the data, made the figures, and wrote the paper. M.Coffin – performed experiments. R.West – designed experiments and analyzed the data. V.M.Fowler – designed the research and study, the experiments and wrote the paper.

## Abbreviations

ED: erythroid differentiation
TMOD: tropomodulin
TPM: tropomyosin
G-actin: globular (monomeric) actin
F-actin: filamentous actin
MEL ds19: mouse erythroleukemia clone ds19
KO: knockout
DMSO: dimethyl sulfoxide solution
HbA: Hemoglobin
CRISPR: clustered regularly interspaced short palindromic repeats
Cas9: CRISPR-associated protein 9
TMB: 3,3’,5,5’-tetramethylbenzidine
FACS: Fluorescence-activated cell sorting
7-AAD: 7-aminoactinomycin D
SSC: side scatter
FSC: forward scatter
FITC: Fluorescein isothiocyanate
PE: phycoerythrin
APC: allophyocyanin
AF488: Alexa-Fluor 488

## Supplemental Figure Legends

**Supplementary Figure S1.**
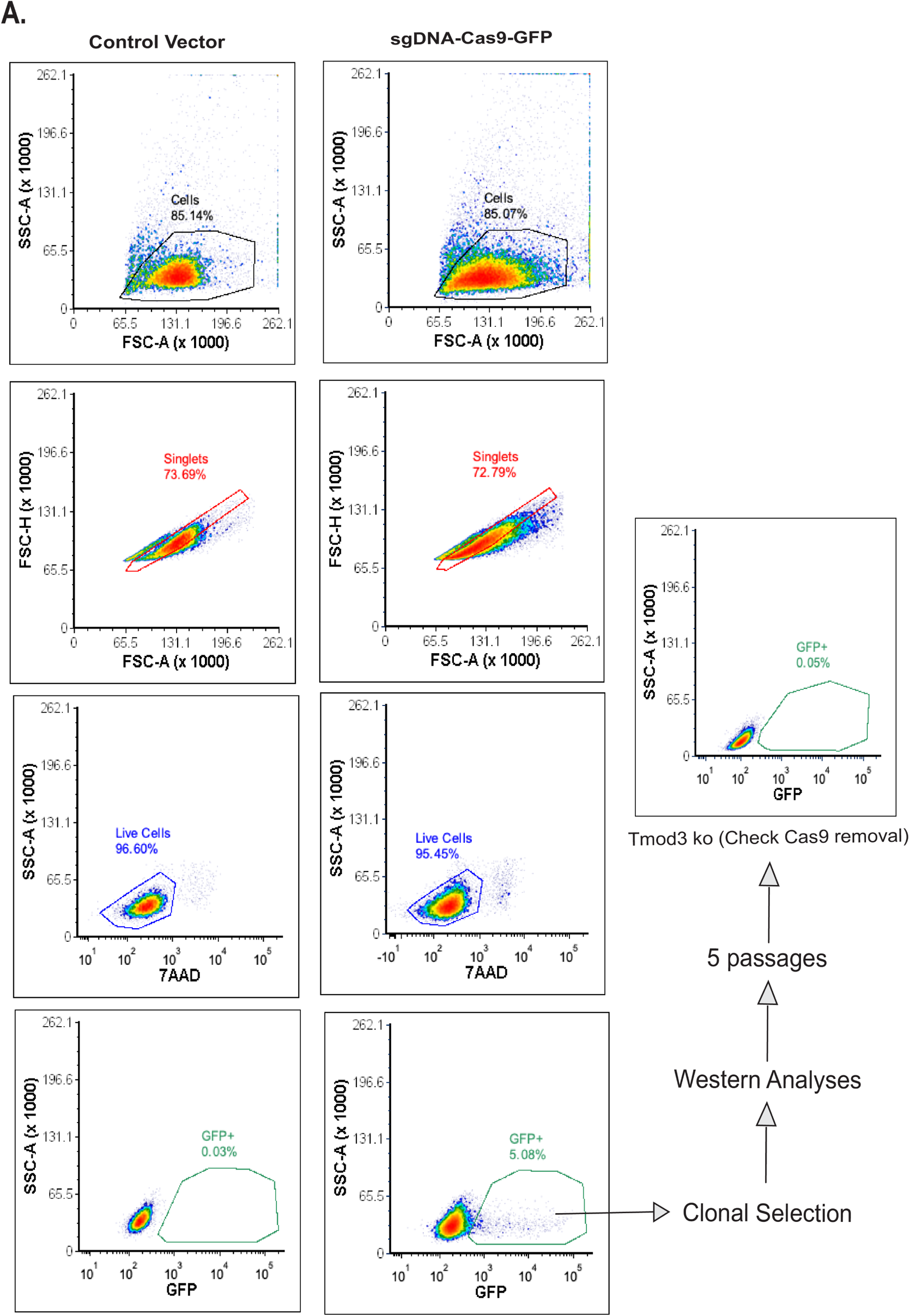
FACS sorting for *Tmod3* knockout cells. (A) Flow cytometry panels showing FACS-sorting for Mel ds19 cells electroporated with plasmid (pSpCas9(BB)-2A-GFP, PX458) expressing sgDNA directed to exon 9b of *Tmod3* gene. A control vector lacking GFP was used to gate for non-GFP fluorescing cells (SSC-A (side scatter-area) vs GFP). The GFP positive cells post-sorting were expanded for clonal selection and screened for loss of Tmod3 via western blotting (data not shown). The selected clone was passaged five times to ensure loss of Cas9-GFP from the cell line and checked via the aforementioned FACS (gating) methodology to ascertain removal of Cas9 from the cells. Cell sorting was carried out on a BD FACSAria Fusion High Speed Cell Sorter.

**Supplementary Figure S2.**
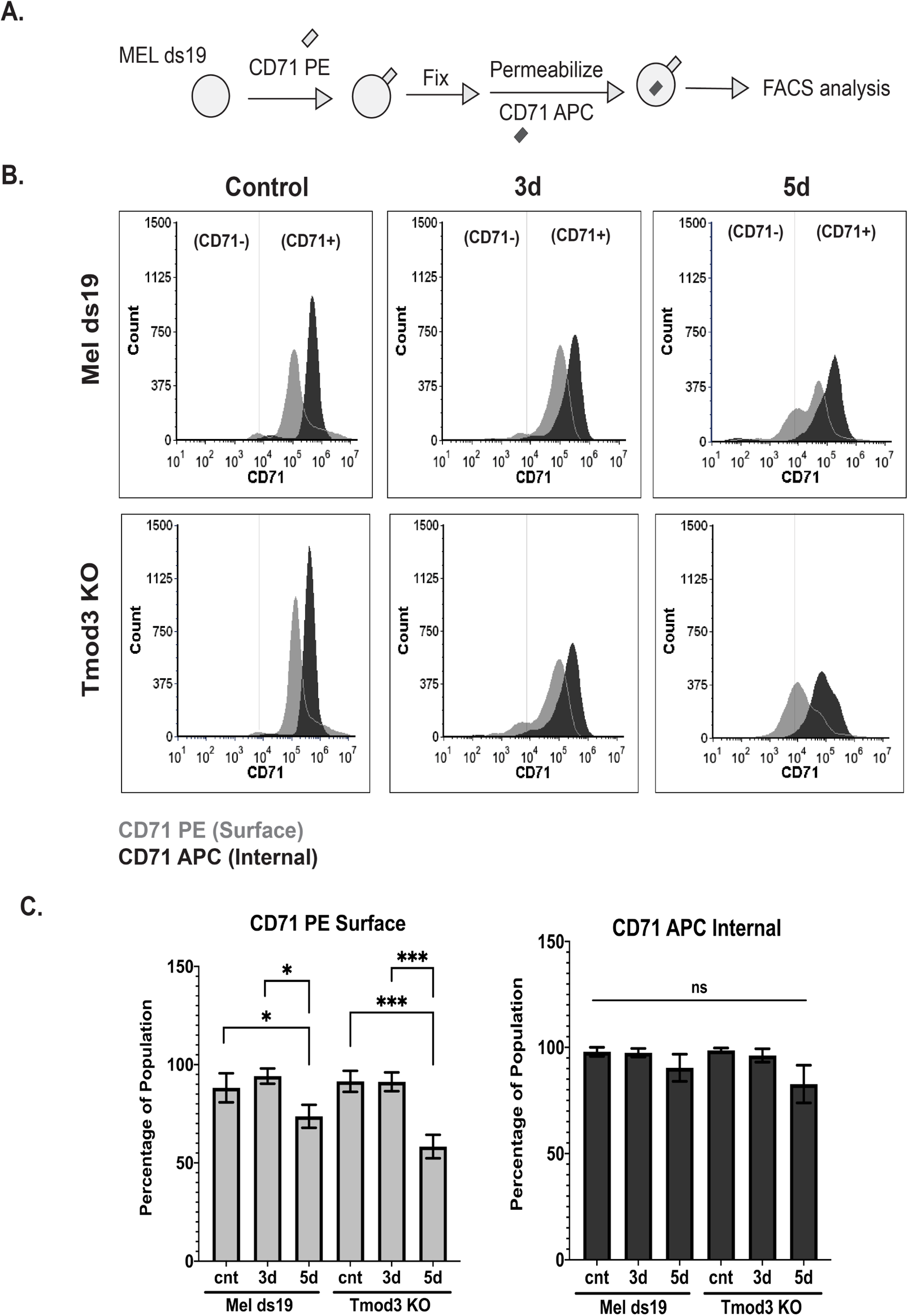
CD71 surface expression during erythroid differentiation is reduced due to *Tmod3* loss. (A) Experimental design for detecting surface and internal CD71 levels via flow cytometry. Cells were stained with anti-CD71 PE antibody followed by fixing and permeabilization using fix and perm kit (according to manufacturer’s instructions, Thermo Fisher), and thereafter labeling the internal CD71 protein with anti-CD71 APC antibody. Cells were subsequently analyzed by flow cytometry using a BD Accuri C6 flow cytometer. (B) Histogram plots for Mel ds19 and *Tmod3* KO cell lines showing surface CD71 PE (light gray) and internal CD71 APC (dark gray) represented as cell count versus maximum fluorescence intensity (MFI). The marker (at ~10^4^ MFI) separates CD71-(unstained) and CD71+ (stained) populations. Histogram plots were generated using FCS express 7 research (Denovo™ Software). (C) Percentage of cells expressing higher than baseline (unstained sample) CD71 fluorescence (MFI) were graphed as a percentage of population for Mel ds19 and *Tmod3* knockout cells at undifferentiated (cnt), 3d and 5d DMSO timepoints. No significant difference was observed when comparing Mel ds19 (5d) vs *Tmod3* KO (5d). Values represent means and error bars ± SD (n=3). Paired student’s T-tests were used to determine significance. * *p* ≤ 0.05, **** *p* ≤ 0.0001.

**Supplementary Figure S3.**
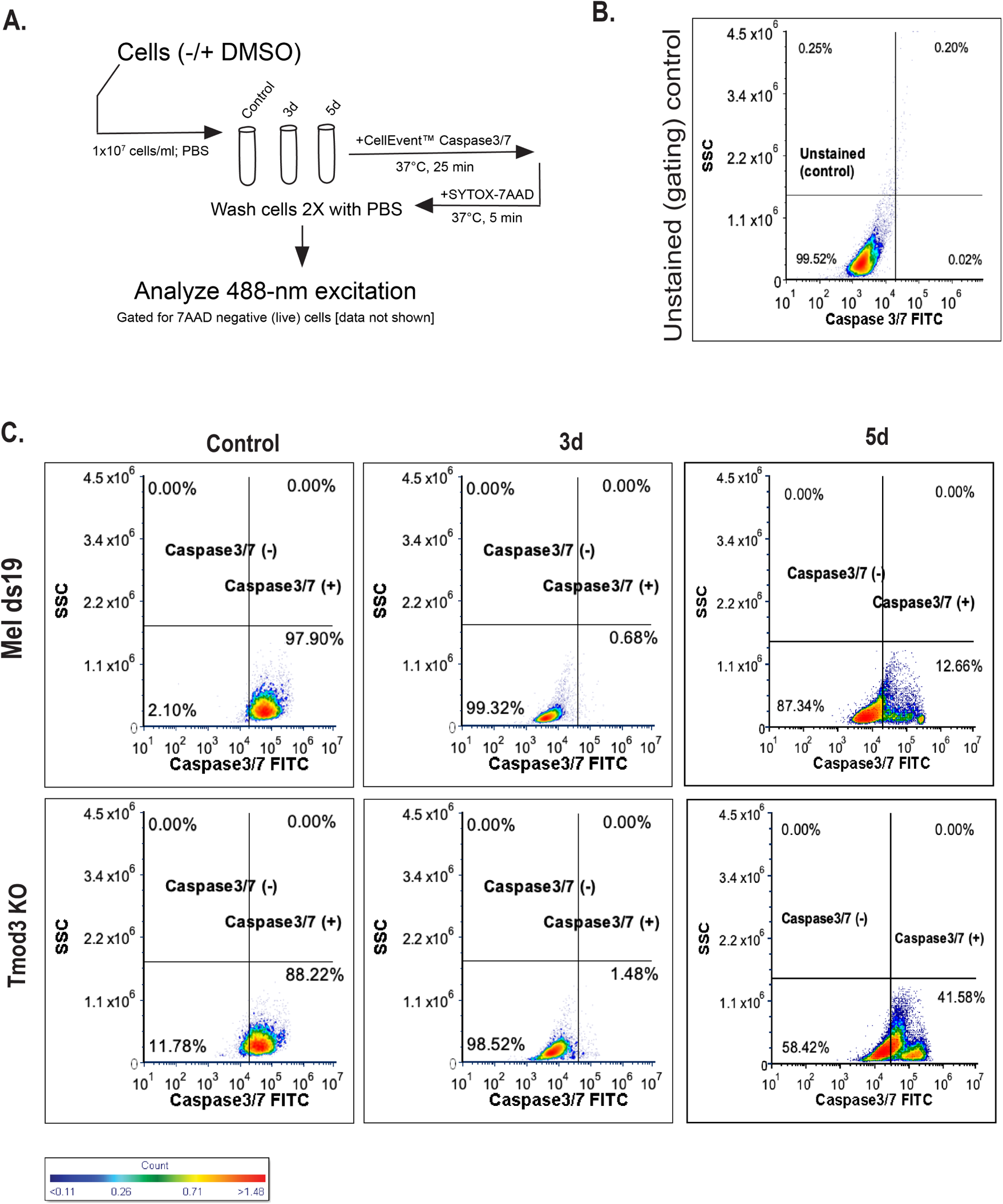
Caspase 3/7 expression is higher in *Tmod3* knockout during erythroid differentiation compared to Mel ds19. (A) Experimental design for Caspase3/7 assay. Cells growing with or without DMSO were washed 2X with PBS before incubating CellEvent™ Caspase3/7 green detection reagent followed by SYTOX-7AAD (as a live/dead stain). Caspase3/7 was detected via excitation with a 488 nm laser in the FITC channel on a BD Accuri C6 flow cytometer. Cells gated for 7AAD negative (live; parent gate, data not shown) were used to gate for the Caspase3/7 positive signal. (B) Unstained Mel ds19 cells were used to establish gating strategy for detection of Caspase3/7 in the FITC channel. A representative flow cytometry panel is shown [SSC(side scatter) vs Caspase 3/7 FITC]. The unstained cell population (99.52%) is denoted in the lower left quadrant for the flow panel. (C) Cells stained with Cell Event™ Caspase 3/7 dye were analyzed via flow cytometry and detected as fluorescence in the FITC channel for Mel ds19 and *Tmod3* knockout cells (undifferentiated/control, 3d and 5d DMSO). Percentage fluorescence intensity is denoted in each gate on lower left (C3/7−) and lower right (C3/7+). The fluorescence intensity (color) denoted as a count legend. All experiments were conducted on BD Accuri C6 flow cytometer and in triplicates (N=3).

**Supplementary Figure S4.**
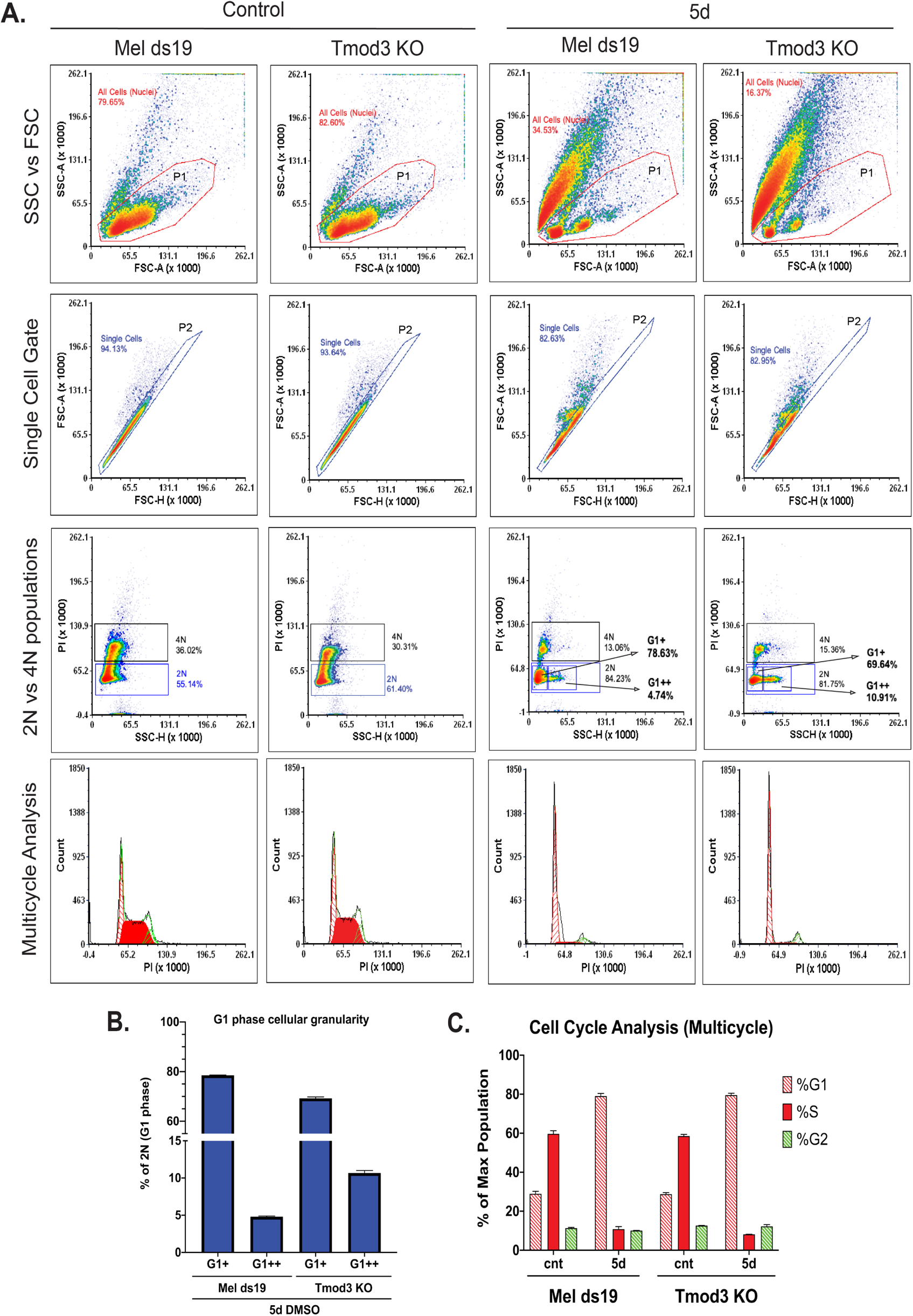
Cell cycle analysis for Mel ds19 and *Tmod3* knockout during erythroid differentiation. (A) Representative flow cytometry panels for Mel ds19 and *Tmod3* knockout cells at control or 5d DMSO timepoints. Cells were stained with propidium iodide (PI) and analyzed on a BD FACS Aria Fusion for cell cycle characterization. Top (row) panels show SSC (side scatter) plotted against FSC (forward scatter) and the gate P1 represents SSC^low^FSC^high^. The next (row) panel – Single Cell Gate represents P1 cells gated for single populations and represented as the P2 gate. In the third row of panels, the 2N vs 4N populations are shown as PI versus SSC-H (side scatter height) with gates 2N and 4N for each subpopulation. For 5d DMSO, Mel ds19 and *Tmod3* KO panels, the 2N population (G1 cells) is sub-gated into two populations – G1+ and G1++ where the latter population indicates cells with greater granularity (internal complexity). The bottom (last row) panels denote histograms for multicycle analysis to model cell cycle stages (%G1, %S, %G2) and are represented as cell counts versus the PI fluorescence. The analysis was conducted on FCS express 7 research (De Novo Software) using the multicycle analysis plugin. (B) For differentiating (5d) Mel ds19 and *Tmod3* knockout cells, the gates on the 2N population (G1+, G1++) were graphed to assess differences in granularity. The percentage of cells in each population was graphed on the Y axes. The *Tmod3* KO G1++ gate indicates that the knockout cells are ~5% more granular than the parental line. (C) Cell cycle parameters (%G1, %S, %G2) from the multicycle analysis generated percent population values for each cell cycle stage. The percentages are shown in the graph with percentage of max population on the Y-axes for control (undifferentiated) and 5d (differentiating) timepoints for respective cell lines. Values represent means, with error bars ± SD (n=3).

**Supplementary Figure S5.**
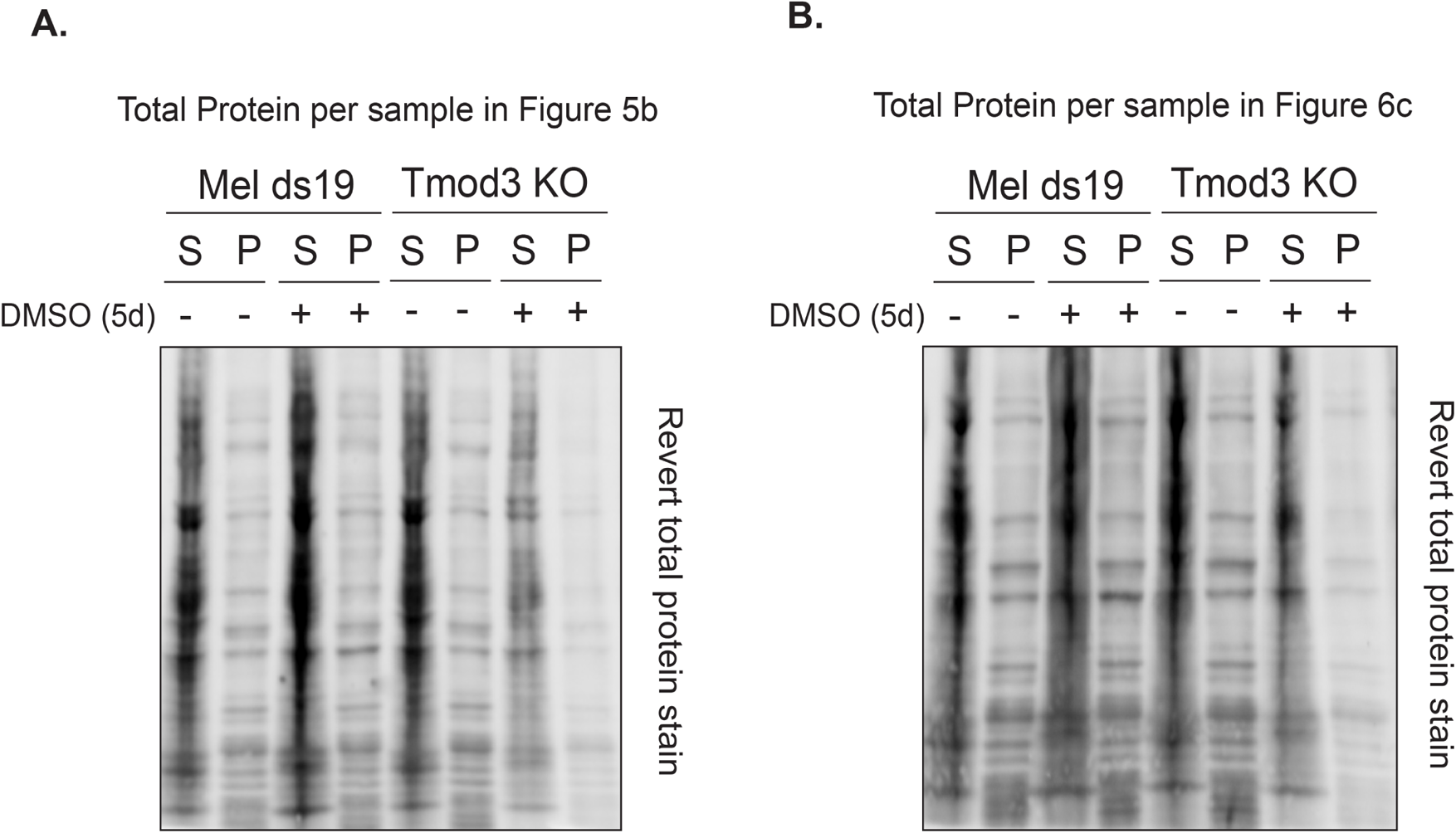
Total protein amounts in S and P fractions in Triton X-100 fractionation assays. Samples from the Triton X-100 fractionation assay denoted as Triton X-100 soluble (S) versus insoluble (P) fractions, were analyzed for total protein prior to blocking and antibody incubation. Post-transfer the PVDF membranes were incubated in Revert™ total protein stain (LiCoR) and visualized in the 700 nm channel using a Biorad ChemiDoc. A representative stained membrane is shown corresponding to samples displayed in figure 5b (A) or figure 6c (B).

---

**Supplemental Table 1.**

| Table S1 |  |  |
| --- | --- | --- |
| Reagent/Resource | Source | Catalog No. |
| Plasmids; Bacterial strains |  |  |
| pSp-Cas9(BB)-2A-GFP (PX458) | Addgene | 48138 |
| DH5α competent cells | Thermo Fisher | 18258012 |
| Plasmid Miniprep Kit | Qiagen | 12943 |
| Chemicals/equipment |  |  |
| DMEM culture media | Fisher Scientific | 10-13-CV |
| Fetal Bovine Serum | GenClone | 25-550H |
| Pen/Strep | Gibco | 15140-122 |
| BbsI | New England Biolabs | R0539S |
| T4 DNA ligase | New England Biolabs | M0202S |
| T4 Polynucleotide kinase | New England Biolabs | M0201S |
| Trypan Blue | Millipore Sigma | 15250061 |
| 7AAD | Biolegend | 420403 |
| Cytofunnel | Thermo Fisher | 59-910-40 |
| Giemsa Stain | Sigma-Aldrich | GS500 |
| TMB Substrate Kit | Pierce (Thermo Fisher) | 34021 |
| BCA Assay Kit | Thermo Fisher | 23250 |
| DMSO | Millipore Sigma | 472301 |
| Annexin V binding buffer (5X) | BioLegend | V13246 |
| Annexin V Brilliant Violet 421 | BioLegend | 640924 |
| CellEvent<sup>TM</sup> Caspase 3/7 | Thermo Fisher | C10423 |
| Propidium Iodide | Sigma-Aldrich | P-4170 |
| IGEPAL CA-630 | Sigma-Aldrich | I8896 |
| Halt<sup>TM</sup> Protease Inhibitor Cocktail | Thermo Fisher | 78429 |
| Halt<sup>TM</sup> Phosphatase Inhibitor Cocktail | Thermo Fisher | 78420 |
| Cytochalasin D | Thermo Fisher | PHZ1063 |
| RNase A | Sigma-Aldrich | R-5000 |
| Triton X-100 | Sigma-Aldrich | X-100 |
| 2X Laemmli Sample Buffer | Biorad | 1610737 |
| Novex Tris-glycine SDS gel (4-12%) | Thermo Fisher | XP04125BOX |
| Novex Tris-glycine SDS gel (4-20%) | Thermo Fisher | XP04205BOX |
| PVDF Immobilon Transfer Membrane | Millipore Sigma | IPFL00010 |
| Fix and Perm Cell Permeabilization kit | Thermo Fisher | GAS003 |
| Revert Total Protein Stain | LiCoR | 926-11011 |
| Antibodies |  |  |
| Tmod3 (Nterm Ab, Chicken Polyclonal, affinity purified) | Genscript | Custom-made |
| Tmod3 (Cterm Ab, Chicken Polyclonal, affinity purified) | Genscript | Custom-made<br>R158181-1 |
| Tmod3 (Mid Ab, Rabbit Polyclonal) | Aviva | ARP55078_P050 |
| Tmod1 (Rabbit polyclonal, affinity purified, R1749 Bleed3A, Tmod3 pre-cleared) | Fowler,1990. J. Cell Biol. | Custom-made |
| Band 4.1R (Rabbit Polyclonal, affinity purified) | Cathy Korsgren & Sam Lux, Harvard Medical School, Boston MA | Gift |
| a1,b1-spectrin (Rabbit polyclonal, affinity purified) | Cathy Korsgren & Sam Lux, Harvard Medical School, Boston, MA | Gift |
| Pan-Actin (C4; Mouse Monoclonal) | EMD Millipore Sigma | MAB1501 |
| Hemoglobin A | Santa Cruz | sc-514378 |
| Tpm1/α-9d (Tpm2,3,5a,5b,6; Sheep Polyclonal) | EMD Millipore Sigma | AB5441 |
| Tpm3.1/γ-9d (5NM1, Sheep Polyclonal) | EMD Millipore Sigma | AB5447 |
| Tpm3.1/Tpm3.2 ((2G10.2; Rabbit Polyclonal) | EMD Millipore Sigma | MABT1335 |
| Tpm4.1/Tm4 (Rabbit Polyclonal) | EMD Millipore Sigma | AB5449 |
| PE Rat Anti-mouse CD71 (Clone C2) | BD Biosciences | 561937 |
| APC Rat Anti-mouse CD71 | Biolegend | 113819 |
| Fc Block | BD Biosciences | 553142 |
| IRDye<sup>®</sup> 800CW Donkey anti-Goat IgG (Secondary Ab) - used with anti-Sheep Primary | LiCoR | 926-32214 |
| IRDye<sup>®</sup> 680RD Donkey anti-Goat IgG (Secondary Ab) - used with anti-Sheep Primary | LiCoR | 926-68074 |
| IRDye<sup>®</sup> 800CW Donkey anti-Rabbit IgG (Secondary Ab) | LiCoR | 926-32213 |
| IRDye<sup>®</sup> 800CW Donkey anti-Mouse IgG (Secondary Ab) | LiCoR | 926-32212 |
| IRDye<sup>®</sup> 680RD Donkey anti-Rabbit (Secondary Ab) | LiCoR | 926-68073 |
| IRDye<sup>®</sup> 680RD Donkey anti-Mouse IgG (Secondary Ab) | LiCoR | 926-68072 |

